# Using deep maxout neural networks to improve the accuracy of function prediction from protein interaction networks

**DOI:** 10.1101/499244

**Authors:** Cen Wan, Domenico Cozzetto, Rui Fa, David T. Jones

## Abstract

Protein-protein interaction network data provides valuable information that infers direct links between genes and their biological roles. This information brings a fundamental hypothesis for protein function prediction that interacting proteins tend to have similar functions. With the help of recently-developed network embedding feature generation methods and deep maxout neural networks, it is possible to extract functional representations that encode direct links between protein-protein interactions information and protein function. Our novel method, STRING2GO, successfully adopts deep maxout neural networks to learn functional representations simultaneously encoding both protein-protein interactions and functional predictive information. The experimental results show that STRING2GO outperforms other network embedding-based prediction methods and one benchmark method adopted in a recent large scale protein function prediction competition.

## Introduction

The realisation of the complex relationships between genotypes and phenotypes has been fostering the collection and analysis of genome-wide datasets of molecular interactions detected from patterns of physical binding, transcript co-expression, mutant phenotypes, etc. Many specialised databases exist to store and integrate such heterogeneous data at different levels of biological complexity. At one end of the scale, the IMEx consortium gathers non-redundant protein-protein interactions (PPIs) from peer-reviewed scientific publications, and provides manually curated details about the experimental conditions [1]. At the opposite end, several resources extend these primary data with indirect or predicted associations to paint a more complete picture for whole organisms [2–5]. For instance, STRING [5] considers experimentally detected PPIs, conserved mRNA co-expression, co-mention in abstracts and papers, interactions from curated databases, conserved gene proximity, gene co-occurrence/co-absence and gene fusion events. Interactions in such databases are typically assigned confidence scores, which can be used for integration purposes [2,6,7]. Not only these data provide valuable direct links between genes and their biological roles, but also form the basis for protein function prediction methods that do not rely on traditional annotation transfers from sequence. Omics data have long offered a suitable opportunity by lending themselves to network representations, where genes or protein products are nodes and edges represent molecular interactions. This modelling approach can be easily exploited using the “guilt-by-association” principle: if the edges reflect biological facts reliably, adjacent nodes have more similar functions than those further away in the network - e.g. because they form a macromolecular complex, or their activities are coordinated in a specific biological process.

The earliest methods therefore transfer annotations from nodes that are either adjacent or within close distance, possibly taking into account the enrichment of the functional labels [8]. Because the network topology is far from uniform and different functions arise from unevenly sized gene sets, using one particular distance or number of neighbours inevitably affects prediction accuracy. More sophisticated algorithms therefore try to group the nodes into functional modules or communities – each associated with a given function – and then make annotation transfers within them [9–14]. The preliminary identification of functionally coherent subgraphs, however, poses additional challenges, which can make module-assisted predictors less accurate than those based on neighbour counting [15]. More recently, network propagation methods have become increasingly popular to address a wide range of problems [16]. They broadcast annotations from labelled proteins to others by running random walks, which visit the nodes in the network randomly until stopping criteria are met [17–19]. If the edges are weighted, this information controls the probability of traversing them; otherwise equal probabilities are used. Because the propagation is affected by node degree and edge weights, this approach reduces the chance of erroneous predictions from highly multifunctional hub proteins to adjacent nodes, which perform fewer functions. Alternatively, the transition probabilities can be used to encode directly the nodes as multi-dimensional features, and thus to make functional annotations with nearest neighbour strategies [20,21]. Cho et al. (2016) [22] and Gligorijević et al. (2018) [23] have instead used them to embed the STRING networks jointly – that is to map nodes to continuous features, which best explain the transition probabilities and the graph topology. The usefulness of the resulting features has been demonstrated for the task of protein function prediction.

This study proposed a novel PPI network-based protein function predicting method, STRING2GO. It adopts deep maxout neural networks to learn a novel type of functional biological network feature representations simultaneously encapsulating both node neighborhoods and co-occurrence functions information. These higher-level representations are learnt in a supervised way by training deep maxout neural networks to output all the terms in biological process domain associated with an input protein - an approach that has led to higher predictive accuracy in the past [24,25]. The experimental results show that STRING2GO significantly outperforms other PPI network embedding-based protein function prediction methods.

## Materials and methods

### Data Collection

Firstly, human proteins were retrieved from the UniProtKB/SwissProt release 2017_05 [26], while the corresponding protein-protein interactions information was retrieved from STRING v10.0 [27] that includes seven component networks from heterogeneous data sources and one integrated network. The mapping between UniProtKB/SwissProt accession numbers and Ensembl protein identifiers adopted in STRING was obtained by using the Biomart tool [28].

Experimentally supported Gene Ontology (GO) term annotations – identified with evidence code EXP, IDA, IPI, IMP, IGI or IEP – were collated from the UniProtKB/SwissProt release 2017_05 and UniProt-GOA release 168 [29], and propagated over “is a” relationships in the Gene Ontology database [30] - GO obo file release 2017-04-28. To assure the feasibility of the following machine-learning experiments, only biological process (BP) annotating at least 100 proteins were initially considered. To guarantee that the predictions are sufficiently specific and informative, this list was subsequently filtered so that only the deepest terms in the ontology were retained − i.e. the terms a and b were kept if and only if there are no “is_a” paths from a to b and from b to a. These steps yielded a vocabulary consisting of 204 BP terms (detailed information is included in Table S1).

The set of human proteins was split into a large subset for GO term-specific classifier training and a small subset for held-out evaluation. 10,667 proteins with at least one cellular component term were initially selected from the whole set. Out of these, 1,000 proteins were randomly selected for held-out evaluation from the subset of well-annotated entries − i.e. those with at least 28, 5 and 14 experimental or electronic biological process, molecular function and cellular component terms respectively. After removing electronic annotations, the held-out set for BP terms contains 982 proteins, while the large set contains 5,000 proteins. In addition, we also create a separated protein-set for a temporal annotation validation by selecting 428 proteins who had no experimental annotation by any 204 BP terms but received at least one after 6 months. The source files were collected from UniProtKB/SwissProt release 2017_11, UniProt-GOA release 174 and GO obo file 2017-10-30.

### Predictive performance evaluation

Predictive performance was evaluated on the ability to annotate both individual labels (GO term-centric) and protein function (protein-centric), following the methodology adopted in [31]. For the GO term-centric evaluation, we calculate the F1 score for evaluating the GO term-specific classifier training quality over 10-fold cross validation on the large training protein-set and the predictive performance on the held-out protein-set. In details, the GO term-centric F1 (i.e. F1_GO_) score is used for evaluating the performance of methods when predicting protein annotations for individual GO terms. As shown in Equation 1, the F1 score is obtained by calculating the harmonic mean of precision and recall values. The precision value (Equation 2) is calculated by dividing the number of true positive (TP) predictions over the summation of true positive and false positive (FP) predictions, while the recall value (Equation 3) is calculated by dividing the number of true positive (TP) predictions over the summation of true positive and false negative (FN) predictions.

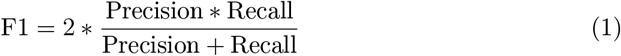

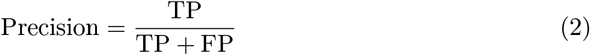

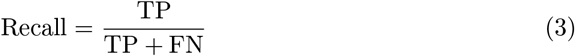

For the protein-centric evaluation, we calculate the F_max_ score by predicting the GO term annotations for the held-out protein-set using the trained GO term-specific classifiers. The F_max_ score is used by CAFA experiments [31] for evaluating the performance of methods when predicting GO term annotations for all protein samples. As shown in Equation 4, the F_max_ score is obtained by choosing the maximum averaged F1 score over all protein samples’ GO term annotation prediction, according to the varied decision threshold. The averaged F1 score for threshold *τ* is calculated by the averaged precision 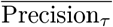 (Equation 5) and recall 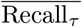 (Equation 6) values. The 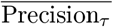 value is calculated by the total amount of precision values for the GO term annotation predictions of all protein sequences *S*, over the number of protein sequence m with at least one GO term annotation predictive posterior probability being equal or greater than the value of threshold *τ*. Analogously, the 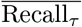 value is calculated by the total amount of recall values for the GO term annotation predictions of all protein sequences *S*, over the total number of protein sequences *n*. Then the corresponding *τ* to Fmax score is used as the prior knowledge to calculate the other type of protein-centric averaged F1 score, i.e. F_*τ*_, for the temporal annotation validation.

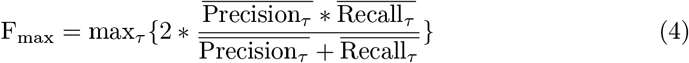

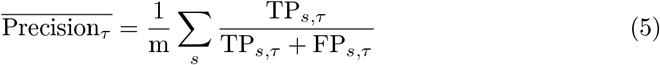

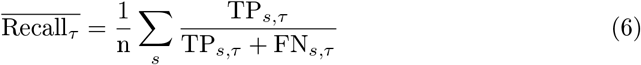

### STRING2GO - a novel protein function prediction method based on learning representations simultaneously encoding the protein-protein interaction and functional annotation information

In general, the STRING2GO method is composed of a three-stage machine learning procedure. As shown in the flow-chart of Fig 1, at the first stage, it adopts the network embedding representation generation methods (e.g. Mashup and Node2vec discussed in this work) to generate the vector representations for individual proteins based on the protein-protein interaction network. Then the Deep Maxout Neural Networks (DMNNs) feed-forward those generated representations as the inputs to a set of GO term annotations of individual proteins as the outputs. The new type of functional representations (denoted as STRING2GO_Embedding_) that simultaneously encode the PPI and protein functional annotation information are extracted from the outputs of the 3^*rd*^ hidden layer of DMNNs after finishing the backward propagation optimisation. Finally, STRING2GO trains a library of Support Vector Machines (SVMs) to predict the posterior probability of annotating individual GO terms to the target proteins. Here, we denote this type of STRING2GO method as STRING2GO_Embedding+SVM_ for clarity. In addition, due to the natural functionality of DMNNs, we also propose another type of STRING2GO method, denoted as STRING2GO_Embedding+gigmoid_, which directly adopts the sigmoid function in the last layer of DMNNs to make predictions.

**Fig 1.**
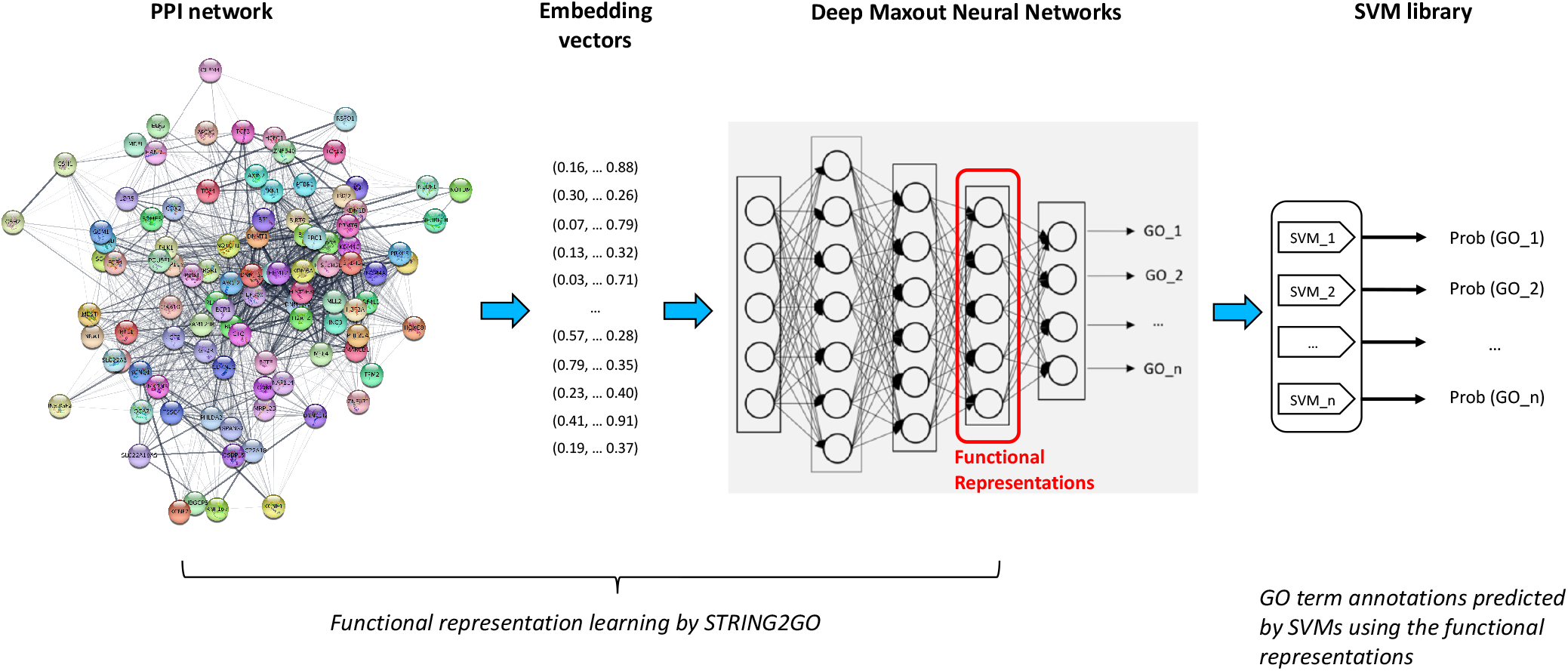
Flow-chart of STRING2GO-based protein function prediction method

In this work, we evaluate the predictive performance of our two types of STRING2GO method on predicting the BP terms located in the deep positions in the GO-DAG, benchmarking with the conventional raw network embedding representations-based method, i.e. Embedding+SVM, that merely adopts the raw network embedding representations to train the SVMs for making predictions.

### Network embedding representation generation

In this work, we adopt two types of network embedding representation generation methods, i.e. Mashup [22] and Node2vec [32], to derive representations from STRING networks. Mashup firstly evaluates the diffusion states of nodes in the network by random walks with a restart approach. Then the truncated singular value decomposition is applied to the diffusion state matrix in order to learn a lower dimensional representation space that optimally approximates the original diffusion states information. The usefulness of the resulting network embedding representations has been demonstrated for a range of functional classification tasks, including function and genetic interaction prediction. As suggested, the best-performing Mashup-derived representations are 800 dimensional and generated by the random-walk sampling strategy with the restart probability of 0.5.

Analogously, Node2vec firstly obtains the node neighborhood information by truncated random walks. Then a Skip-gram [33,34] shallow neural network is used to generate a representation space, where the nodes contain the maximum likelihood of preserving corresponding node neighborhood information. In this work, the neighborhood information was sampled through random walks of length ten, which were biased towards close neighbors by setting the parameter *q* to 2. We also evaluate the performance of representations in different dimensions, i.e. 32, 64, 128, 256 and 512, generated from all different STRING networks [20,21].

### Deep maxout neural networks training

Deep Maxout Neural Networks (DMNNs) are used for learning the more abstract representations simultaneously encoding the PPI network information and the patterns of term co-occurrence in the biological process functional domain. The network architecture was implemented using the Keras package with Theano backend and consisted of three fully connected hidden layers, followed by an output layer with as many neurons as the numbers of terms selected for the biological process functional domain. Each hidden layer had batch-normalized inputs [35], which were combined through maxout units [36], and were subject to dropout [37] in the course of training. A sigmoid function was used to activate the output neurons.

To limit the computational requirements for model optimization, the initial 10-fold cross validation (with random split of instances) experiments were run in order to identify the best combination of optimizer (AdaGrad), number of maxout units (3), learning rate (0.05), batch size (100 elements), and number of epochs (150), keeping fixed the weight initialisation (Glorot uniform method) and the number of units in all hidden layers, by considering the highest F1_GO_ scores for predicting all 204 BP terms. Subsequent training stages were aimed at selecting the optimal dimensions of hidden layers that lead to the highest median F1_GO_ scores (here rounded to two decimal places), from a limited set of options (300, 500, 700 and 1,000). In addition, we also evaluate the predictive performance when using the same dimensions for both input features and the 3^*rd*^ hidden layer outputs. Note that, due to the well-known curse of dimensionality issue [38], if more than two different dimensions of the 3^*rd*^ hidden layer outputs obtain the same median F1_GO_ scores, we only choose the lowest ones as the optimal dimensions.

### Support vector machine training

Scikit-learn [39] was used to train a set of GO term-specific Support Vector Machines (SVMs) with a radial basis function (RBF) kernel, the parameters of which were identified through a grid search as those maximising the F1_GO_ score across the stratified 10-fold cross validation experiments. To train each classifier, the set of positive instances consisted of the proteins annotated with the target GO term *t* or its descendants, while the set of negative instances are all remaining proteins not annotated with the target GO term or its descendants. Finally, the well-known Platt scaling method [40] was used to transform the predictive scores of individual SVMs into a probability distribution of binary classes. The data and code can be accessed via https://github.com/psipred/STRING2GO

## Results

We firstly report the experimental results about evaluating the predictive information included in different STRING networks that are used for generating the raw network embedding representations by two different methods, i.e. Mashup and Node2vec. Then we evaluate the predictive performance of the STRING2GO-learnt functional representation (i.e. STRING2GO_Mashup_ and STRING2GO_Node2vec_) by comparing with their corresponding raw network embedding representations. We also compare the performance of Mashup and Node2vec methods when they are used to generate the raw network embedding representations or be the component methods of STRING2GO to learn the functional representations. Finally, we further compare all prediction methods involved in this work, also benchmarking with the Naïve method [31].

### Predictive power included in different STRING networks

To begin with, we compare the predictive power of different STRING networks by adopting the Mashup or Node2vec-generated network embedding representations as the inputs of DMNNs for predicting protein function (i.e. STRING2GO_Mashup+sigmoid_ and STRING2GO_Node2vec+sigmoid_). Overall, the Combinedscore network-derived embedding representations show the best predictive performance among all different STRING networks-derived ones when using either Mashup or Node2vec methods, while the Textmining network-derived representations also obtain the competitive predictive accuracy. As shown in the 4th and 7th columns of Table 1, the Combinedscore network-derived representations obtain the highest median F1_GO_ (hereafter, denoted by 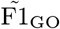) scores (0.23 and 0.17) using Mashup and Node2vec respectively. The Combinedscore network also contains the largest number of proteins, interactions and the highest coverage (as shown in the columns 8-10 of Table 1), when mapping the STRING network-included proteins to the training protein-set. The Textmining network-derived representations obtain the second highest 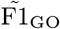 score (0.22) using the Mashup method, while also obtain the same highest 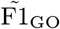 score (0.17) using the Node2vec method. Moreover, in terms of the predictive information included in other component networks, the Experimental network-derived embedding representations show the third highest predictive accuracy, since they obtain sequentially higher 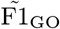 scores than the ones derived by the Database and Coexpression networks respectively. Note that, the embedding representations derived from Neighbourhood, Cooccurrence and Fusion networks show poor predictive performance, since their 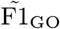 scores are all equal to zero, and the mapping coverages are all lower than 21.0%. Hereafter, we consider learning the functional representations by STRING2GO only from those 5 networks including relatively rich PPI information and high coverage.

**Table 1.** The optimal dimensions of raw network embedding representations and the corresponding 3^*rd*^ hidden layer outputs (a.k.a. the STRING2GO-learnt functional representations) with their corresponding predictive power for biological process terms prediction, and the main characteristics of different STRING networks

| STRING Networks | Mashup |  |  | Node2vec |  |  | No. Proteins | No. Interactions | Coverage on Training set |
| --- | --- | --- | --- | --- | --- | --- | --- | --- | --- |
| | Input | $3^{rd}$ Hidden | $\tilde{F1}_{GO}$ | Input | $3^{rd}$ Hidden | $\tilde{F1}_{GO}$ | | | |
| Combinedscore | 800 | 800 | **0.23** | 128 | 500 | **0.17** | **19247** | **8548002** | **93.4%** |
| Textmining | 800 | 700 | 0.22 | 128 | 1000 | **0.17** | 19088 | 7632934 | 93.3% |
| Experimental | 800 | 700 | 0.19 | 128 | 1000 | 0.13 | 16858 | 3473862 | 90.4% |
| Coexpression | 800 | 700 | 0.14 | 256 | 700 | 0.09 | 12774 | 1537924 | 72.0% |
| Database | 800 | 700 | 0.11 | 128 | 700 | 0.04 | 7937 | 424860 | 56.9% |
| Neighborhood* | 800 | 300 | 0.00 | 32 | 32 | 0.00 | 3514 | 152248 | 20.9% |
| Cooccurrence* | 800 | 300 | 0.00 | 32 | 32 | 0.00 | 2754 | 47478 | 16.6% |
| Fusion* | 800 | 300 | 0.00 | 32 | 32 | 0.00 | 1495 | 4120 | 9.7% |

\* : Note that those STRING networks obtain 0.00 of $\tilde{F1}_{GO}$ scores with all different dimensions, only the lowest dimensions are reported.

We then report the optimal dimensions of network embedding representations derived by Mashup and Node2vec methods from those 5 STRING networks. According to the suggestion in [22], we define 800 as the optimal dimensions for the input network embedding representations derived by Mashup. In terms of the Node2vec-derived network embedding representations, as shown in the 5th column of Table 1, 128 are the overall optimal dimensions, since 4 out of 5 network-derived embedding representations in 128 dimensions obtain the highest 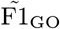 scores for predicting 204 biological process terms. We then report the optimal dimensions of the STRING2GO-learnt functional representations (a.k.a. the 3^*rd*^ hidden layer outputs of DMNNs) w.r.t. the corresponding optimal dimensions of raw network embedding representation inputs. Generally, STRING2GO encodes the functional predictive information in a high dimensional representation space (ranging from 500 – 1000 dimensions), when using either Mashup or Node2vec as the raw network embedding representation generation method. As shown in the 3^*rd*^ and 6^*th*^ columns of Table 1, the optimal dimensions of the 3^*rd*^ hidden layer outputs vary between 500 to 1000. Recall that we also evaluate the cases when the dimensions of the 3^*rd*^ hidden layer outputs are the same to the dimensions of raw network embedding representation inputs. None of the functional representations based on Node2vec-derived network embedding representations obtain higher 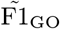 scores when using the same dimensions of inputs as the dimensions of 3^*rd*^ hidden layer outputs, e.g. using 128 as the dimensions of both representation inputs and the 3^*rd*^ hidden layer outputs.

### The functional representations learnt by STRING2GO encode higher predictive power than the corresponding raw network embedding representations

We evaluate the predictive performance of STRING2GO-learnt functional representations by conducting pairwise comparisons with the corresponding raw network embedding representations respectively. Generally, in terms of GO term and protein-centric metrics, both STRING2GO_Mashup_ and STRING2GO_Node2vec_ functional representations obtain higher predictive accuracy than Mashup and Node2vec-derived raw network embedding representations. In detail, during the GO term-specific classifier training stage, as shown in Fig 2.a-2.e, both orange and green bars are lower than other ones. This fact indicates better classifier training quality by using STRING2GO_Mashup+SVM_, STRING2GO_Node2vec+SVM_, STRING2GO_Mashup+Sigmoid_ and STRING2GO_Node2vec+Sigmoid_ than the ones obtained by Mashup+SVM and Node2vec+SVM, when using all five different STRING networks to generate embedding representations.

**Fig 2.**
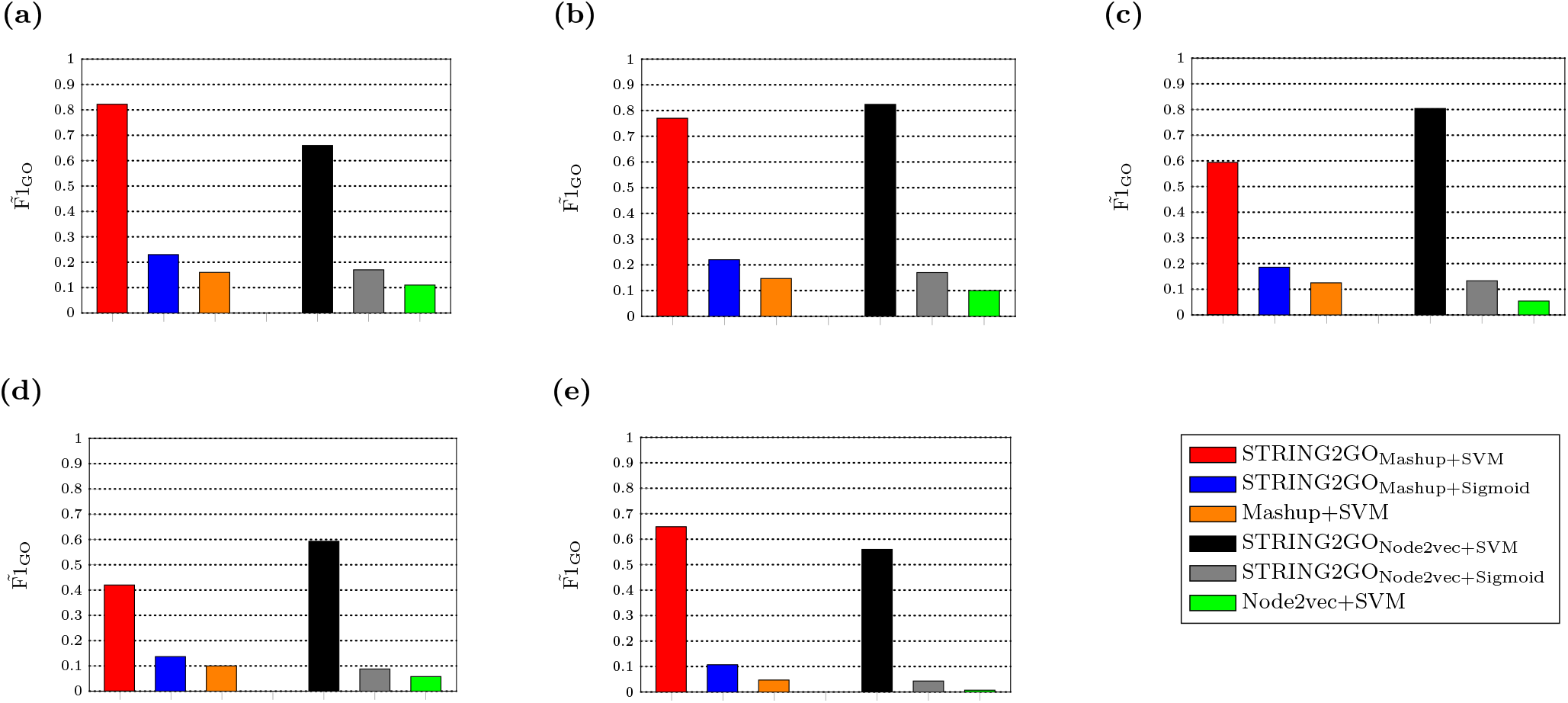
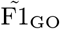 scores obtained by network embedding representations and the corresponding STRING2GO-learnt functional representations based on (a) Combinedscore, (b) Textmining, (c) Experimental, (d) Database and (e) Coexpression networks by using SVM or Sigmoid function over the 10-fold cross validation during the GO term-specific classifiers training stage

The held-out evaluation results further confirm that the STRING2GO-learnt functional representations contain higher predictive information. As shown in Table 2, the 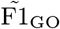 scores obtained by STRING2GO_Mashup+SVM_ and STRING2GO_Node2vec+SVM_ reach to 0.270 and 0.182 respectively, whereas the 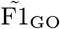 scores obtained by Mashup+SVM and Node2vec+SVM are both equal to 0.000. This pattern is consistent when adopting all other types of STRING component networks, except STRING2GO_Node2vec+SVM_ and Node2vec+SVM both obtain zero 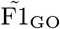 scores when using the Coexpression network to generate the raw embedding representations (as shown in Table 2). STRING2GO_Mashup+Sigmoid_ and STRING2GO_Node2vec+Sigmoid_ also respectively obtain higher 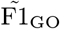 scores than Mashup+SVM and Node2vec+SVM based on all five different STRING networks. The scatter-plots in Fig 3 show the pairwise comparisons of F1_GO_ scores obtained by different methods, and the dashed-lines indicate the median values of difference between pairs of F1_GO_ scores. In detail, Fig 3.a-3.d show that almost all dots (in blue) drop in the area above the diagonal, indicating higher F1_GO_ scores for predicting the majority of BP terms by using the functional representations learnt by STRING2GO based on the Combinedscore network by using either SVM or Sigmoid function as the classification algorithm. As shown in Fig 3.e-3.t This pattern is consistently observed when applying on almost all other four different STRING networks, except the Coexpression network that leads to competitive performance between STRING2GO_Node2vec_ and Node2vec, since the dashed-lines in Fig 3.s and Fig 3.t are almost overlapping on the diagonal. The Wilcoxon signed-rank test results in Table S3 further confirm that the STRING2GO-learnt functional representations obtain significantly higher GO term-centric F1_GO_ scores than the raw network embedding representations.

**Table 2.** Summary on experimental results obtained by different network embedding representations and corresponding functional representations based on Combinedscore, Textmining, Experimental, Database and Coexpression networks working with different classification algorithms during held-out evaluation and temporal annotation validation

| Prediction Methods | Combinedscore |  |  | Textmining |  |  | Experimental |  |  | Database |  |  | Coexpression |  |  |
| --- | --- | --- | --- | --- | --- | --- | --- | --- | --- | --- | --- | --- | --- | --- | --- |
|  | Held-out |  | Temporal | Held-out |  | Temporal | Held-out |  | Temporal | Held-out |  | Temporal | Held-out |  | Temporal |
| | $\tilde{F}_{1GO}$ | $F_{max}$ | $F_\tau$ | $\tilde{F}_{1GO}$ | $F_{max}$ | $F_\tau$ | $\tilde{F}_{1GO}$ | $F_{max}$ | $F_\tau$ | $\tilde{F}_{1GO}$ | $F_{max}$ | $F_\tau$ | $\tilde{F}_{1GO}$ | $F_{max}$ | $F_\tau$ |
| *Mashup-based* |  |  |  |  |  |  |  |  |  |  |  |  |  |  |  |
| STRING2GO<sub>Mashup</sub>+SVM | **0.270** | **0.497** | 0.309 | **0.275** | **0.483** | 0.296 | 0.146 | **0.450** | **0.263** | 0.130 | 0.412 | 0.225 | 0.116 | **0.392** | **0.258** |
| STRING2GO<sub>Mashup</sub>+Sigmoid | 0.237 | 0.495 | 0.312 | 0.239 | 0.478 | 0.290 | **0.183** | 0.442 | 0.247 | **0.131** | **0.427** | 0.144 | **0.121** | **0.392** | 0.247 |
| Mashup+SVM | 0.000 | 0.470 | 0.290 | 0.000 | 0.463 | 0.287 | 0.000 | 0.420 | 0.229 | 0.000 | 0.392 | **0.238** | 0.000 | 0.371 | 0.242 |
| *Node2vec-based* |  |  |  |  |  |  |  |  |  |  |  |  |  |  |  |
| STRING2GO<sub>Node2vec</sub>+SVM | 0.182 | 0.458 | **0.319** | 0.115 | 0.446 | 0.290 | 0.124 | 0.422 | 0.256 | 0.087 | 0.353 | 0.169 | 0.000 | 0.349 | 0.236 |
| STRING2GO<sub>Node2vec</sub>+Sigmoid | 0.187 | 0.471 | 0.312 | 0.188 | 0.472 | **0.314** | 0.143 | 0.440 | 0.258 | 0.111 | 0.408 | **0.238** | 0.043 | 0.381 | 0.246 |
| Node2vec+SVM | 0.000 | 0.444 | 0.293 | 0.000 | 0.437 | 0.278 | 0.000 | 0.418 | 0.249 | 0.000 | 0.386 | 0.221 | 0.000 | 0.360 | 0.219 |
| * Naive | N/A | 0.363 | 0.254 |  |  |  |  |  |  |  |  |  |  |  |  |

**Fig 3.**
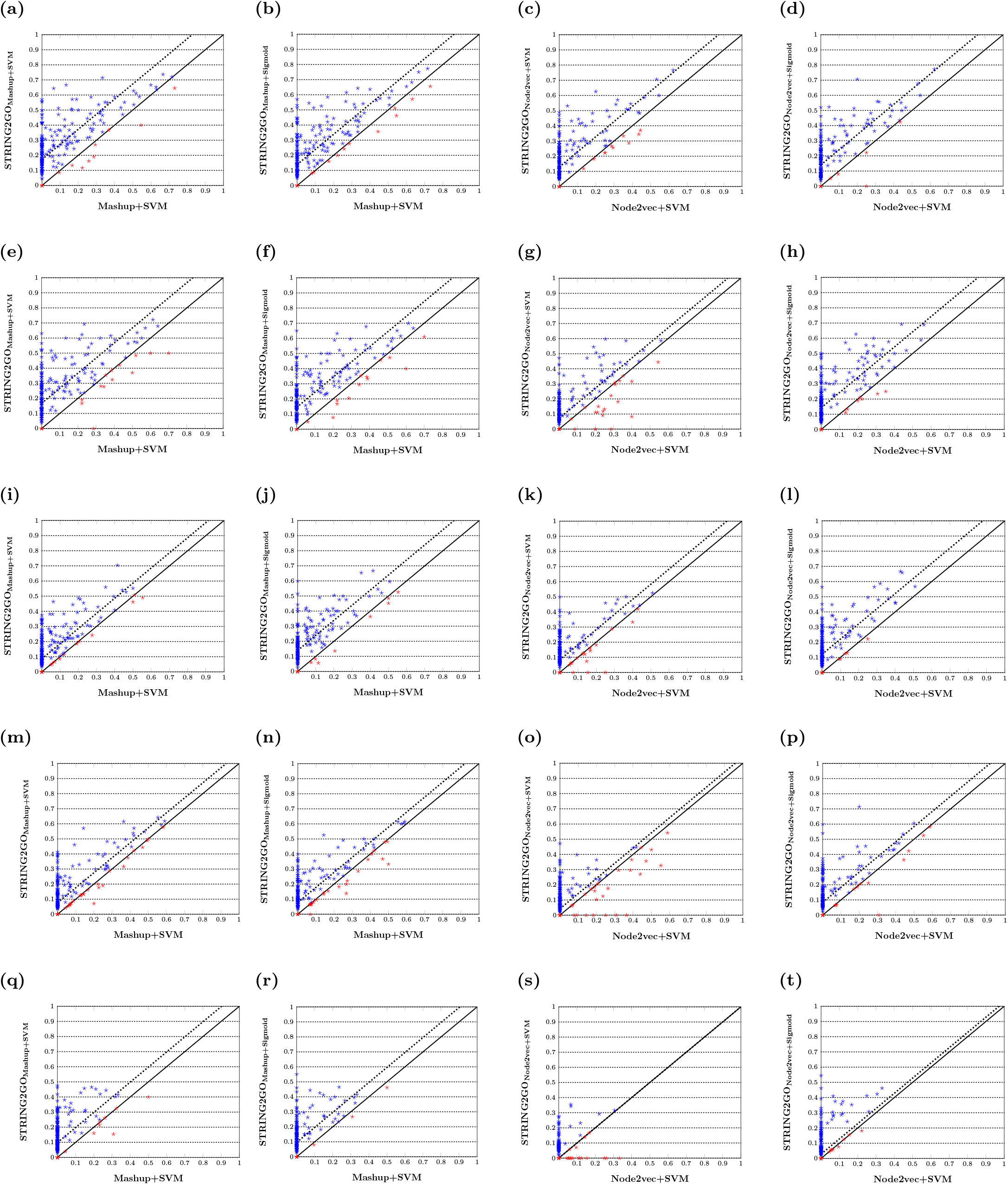
F1_GO_ scores obtained by different network embedding representations and the corresponding STRING2GO-learnt functional representations based on (a-d) Combinedscore, (e-h) Textmining, (i-l) Experimental, (m-p) Database and (q-t) Coexpression networks by using SVM or Sigmoid function for classification

From the perspective of protein-centric evaluation (i.e. considering the F_max_ and F_*τ*_ metrics), the STRING2GO-learnt functional representations also obtain higher predictive accuracy based on the Combinedscore network. As shown in Table 2, the functional representations STRING2GO_Mashup_ and STRING2GO_Node2vec_ both obtain higher F_max_ scores (i.e. 0.497 and 0.458 obtained by using SVM, 0.495 and 0.471 obtained by using Sigmoid function) than the network embedding representations generated by Mashup and Node2vec (i.e. 0.470 and 0.444 obtained by using SVM). The precision-recall curves in Fig 4.a also show that the STRING2GO-learnt functional representations obtain higher precision and recall values simultaneously, since the middle parts of red and blue curves locate in higher position than the orange one, while the middle parts of grey and black curves also locate in higher position than the green one. As shown in Table 2 and Fig 4.b-4.e, this pattern is consistent when adopting the other four types of STRING component networks to generate representations, except STRING2GO_Node2vec+SVM_ obtaining lower F_max_ scores than Node2vec+SVM based on the Database and Coexpression networks.

**Fig 4.**
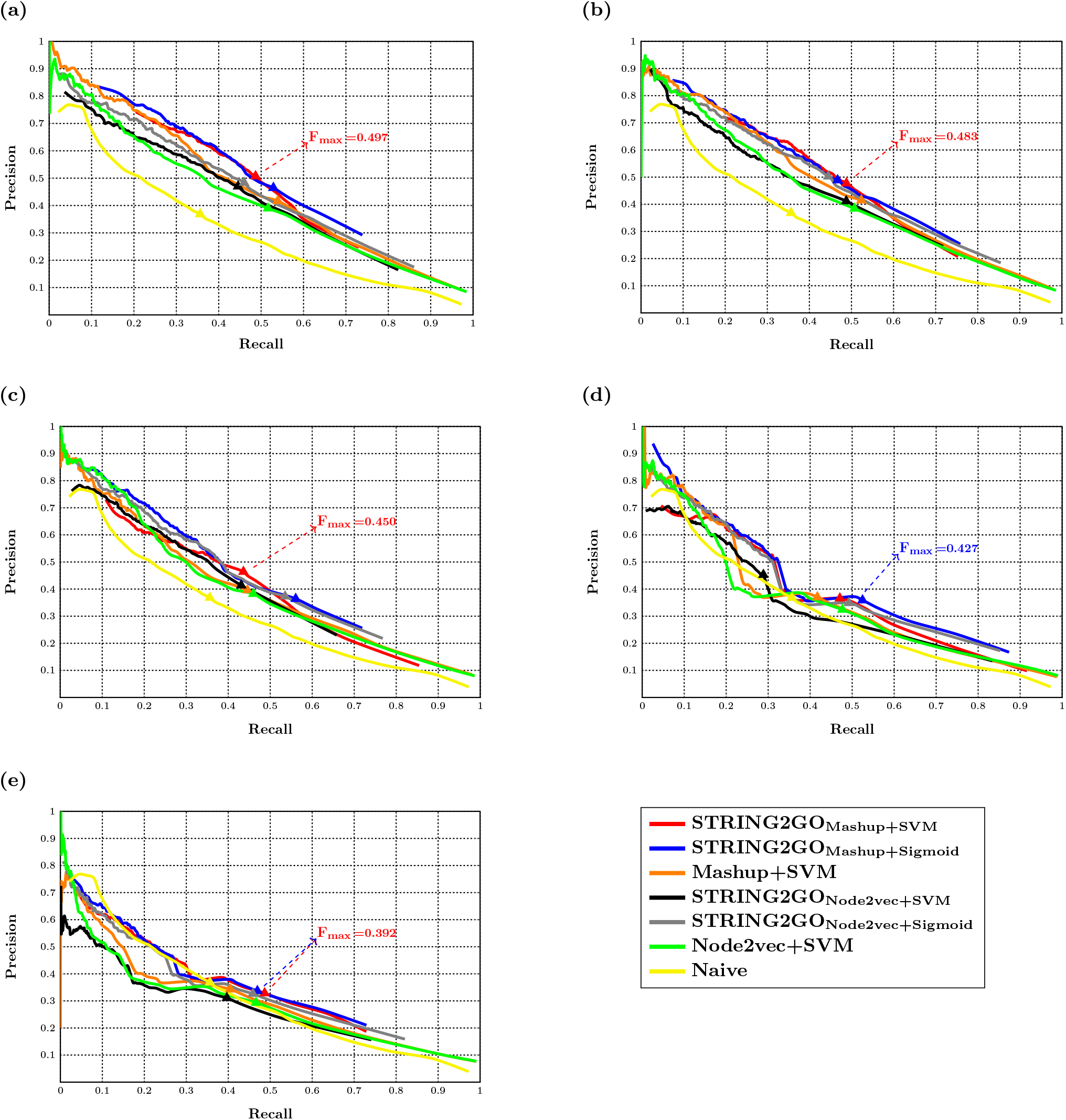
Precision-recall curves of different methods and the Fmax scores obtained by the best-performing methods based on (a) Combinedscore, (b) Textmining, (c) Experimental, (d) Database and (e) Coexpression networks

Analogously, the functional representations STRING2GO_Mashup_ and STRING2GO_Node2vec_ obtain higher F_*τ*_ scores based on the Combinedscroe network (0.309 and 0.319 obtained by SVM, while 0.312 obtained by Sigmoid function) than the raw network embedding representations generated by Mashup and Node2vec (0.290 and 0.293 by using SVM). This pattern is consistent when using all other STRING networks, except the Database network which only leads to higher F_*τ*_ score obtained by STRING2GO_Node2vec+Sigmoid_ than the one obtained by Node2vec+SVM.

### The raw network embedding representations derived by Mashup show higher predictive power

We also compare the predictive performance of Mashup and Node2vec-derived network embedding representations and the corresponding STRING2GO-learnt functional representations respectively. Generally, the raw network embedding representations derived by Mashup and Node2vec methods obtain competitive predictive accuracy by using SVM as the classification algorithm. To begin with, during the training stage, the 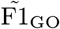 score obtained Mashup+SVM is higher than the one obtained by Node2vec+SVM based on the Combinedscore network, since the orange bar is higher than the green one in Fig 2.a. However, both Mashup+SVM and Node2vec+SVM obtain poor predictive performance on the held-out evaluation, due to the zero 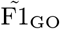 scores. But the statistical significance test results (see Table S2) show that the former still outperforms the latter. Those patterns are consistent when using all other 4 types of STRING networks to generate the raw embedding representations, as reported in Fig 2.b-2.e, Tables 2 and S1. In terms of the protein-centric evaluation, Mashup+SVM obtains a higher F_max_ score (0.470) than Node2vec+SVM (0.444). The Combinedscore network-based precision-recall curves in Fig 4.a confirm that the orange curve locates in higher position than the green one. Those patterns are also consistent in cases when using other four different STRING component networks to generate representations, as shown in Fig 4.b–4.e. However, Node2vec+SVM outperforms Mashup+SVM on the temporal annotation validation. As reported in Table 2, although the latter obtains higher F_*τ*_ score based on three STRING component networks (i.e. Textmining, Database and Coexpression), the former obtains the highest F_*τ*_ score (0.293) based on the Combinedscore network.

We then further conduct comparisons on predictive performance of two different STRING2GO-learnt functional representations respectively based on Mashup and Node2vec-derived raw network embedding representations. During the GO term-specific classifiers training stage, STRING2GO_Mashup_ obtains higher 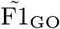 scores than STRING2GO_Node2vec_ by using either SVM or Sigmoid function as the classification algorithm, based on the Combinedscore and Coexpression networks. As shown in Fig 2.a and 2.e, where red and blue bars are higher than the black and grey ones respectively. When using the other 3 STRING component networks, STRING2GO_Node2vec_ obtains higher 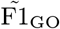 scores by using SVMs, whereas STRING2GO_Mashup_ still outperforms the former by using Sigmoid function as the classification algorithm.

The held-out evaluation results in Table 2 show a consistent pattern that STRING2GO_Mashup_ obtains higher 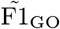 scores (statistically significant according to Table S2) and F_max_ scores than STRING2GO_Node2vec_ based on the Combinedscore network by using either SVM or Sigmoid function, respectively. As shown in Fig 4.a, the majority parts of the red and blue curves clearly locate in higher position than the black and grey ones. Those patterns are consistent when using the other 4 STRING networks, as shown in Table 2 and Fig 4.b-4.e. However, STRING2GO_Node2vec_ obtains better predictive performance during the temporal annotation validation, since the former obtains the highest F_*τ*_ score (0.319) by using SVM (based on the Combinedscore network) among all methods when adopting all different STRING networks.

### The STRING2GO-learnt functional representations with support vector machines obtain the highest accuracy on predicting 204 BP terms

We then compare all prediction methods discussed in previous sections, i.e. two types of STRING2GO methods (i.e. STRING2GO_Embedding+SVM_ and STRING2GO_Embedding+Sigmoid_) adopting two types of raw network embedding representations (i.e. the ones generated by Mashup and Node2vec respectively), and the methods that only exploit the raw network embedding representations to make predictions by using SVM as the classification algorithm. We also compare those methods with the Naïve prediction method [31], which makes predictions by considering the annotation frequency in the database as the prior knowledge. Overall, STRING2GO_Embedding+SVM_ is the best-performing method according to both the GO term and protein-centric metrics. During the GO term-specific classifiers training stage, STRING2GO_Mashup+SVM_ and STRING2GO_Node2vec+SVM_ obtain almost the same highest 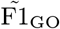 scores among all prediction methods by using all different STRING networks. As shown in Fig 2, the latter obtains the highest GO score (0.824) based on the Textmining network, while the former obtained almost the same highest 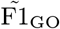 score (0.822) based on the Combinedscore network. The held-out evaluation results also confirm that STRING2GO_Mashup+SVM_ obtains the highest 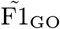 score (0.275) by using the Textmining network, while also obtains the significantly higher F1_GO_ scores than other methods basing on the Combinescore network (see Friedman test with Holm *post-hoc* correction results in Table S3). STRING2GO_Mashup+SVM_ obtains the highest F_max_ score (0.497) based on the Combinedscore network and higher F_max_ scores than all other methods based on all other STRING networks except the Database network. In terms of the F_*τ*_ score metric, STRING2GO_Node2vec+SVM_ obtains the highest F_*τ*_ score (0.319) by using the Combinedscore network among all methods based on all different STRING networks.

The second best performing method is STRING2GO_Embedding+Sigmoid_. STRING2GO_Mashup+Sigmoid_ obtains higher 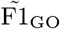 scores than either Mashup+SVM or Node2vec+SVM during the classifier training stage. It also obtains the second highest 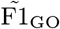 scores during the held-out evaluation based on 2 out of 5 networks (except the case when STRING2GO_Mashup+Sigmoid_ obtains the highest 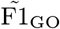 score based on the Experimental, Database and Coexpression networks). From the perspective of protein-centric metrics, STRING2GO_Mashup+Sigmoid_ obtains the second highest F_max_ based on 3 out of 5 STRING networks, while STRING2GO_Node2vec+Sigmoid_ obtains the overall second highest F_*τ*_ score (0.314) based on the Textmining network.

In addition, all of those methods discussed above obtains higher F_max_ scores than the Naïve prediction method based on almost all 5 individual STRING networks (as the yellow curves shown in Fig 4.a-4.e), with exception of STRING2GO_Node2vec+SVM_ based on the Database and Coexpression networks and Node2vec+SVM based on the Coexpression network. All those methods also obtain higher F_*τ*_ scores than the Naïve prediction method based on the Combinedscore and Textmining networks.

## Discussion

Overall, as discussed in previous sections, the functional representations learnt by STRING2GO show substantial improvement on the predictive power of the raw network embedding representations. We further investigate the improvement of predictive power of the STRING2GO-learnt functional representations by evaluating the enlarged distances between two classes of training protein samples. We firstly calculate the Euclidean distance between the centroids of two classes by using the Mashup-based representations’ values standardized into the range of (0,1) in the same dimensional space, i.e. 800 dimensions for both Mashup and STRING2GO_Mashup_. Then we calculate the correlation coefficient between the distances and F1_GO_ scores obtained by held-out evaluation. As shown in Fig 5.a, the *x* axis denotes the distance between two classes calculated by using either the raw Mashup-derived network embedding representations (blue), or the corresponding functional representations (red) STRING2GO_Mashup_, based on the Combinedscore network, while the *y* axis denotes the corresponding F1_GO_ score obtained by adopting those different representations working with SVMs to predict individual BP terms. It is obvious that the distances between two classes of proteins for individual GO terms are all enlarged by STRING2GO, while the correlation coefficient values between distances and F1_GO_ scores for both types of representations are positive, indicating that the larger distances lead to higher predictive accuracy.

**Fig 5.**
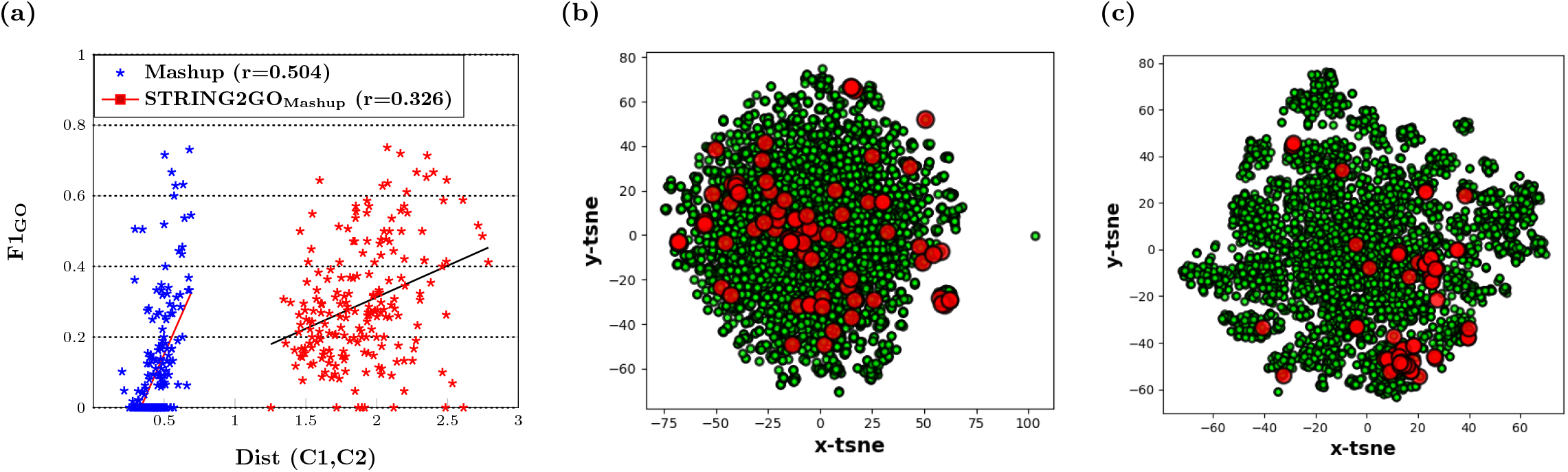
(a) Linear relationship between distances of two classes protein samples and F1_GO_ scores obtained by Mashup-derived network embedding representations and the corresponding functional representations on classifier training stage (c) The 2D space visualization of distribution of protein samples belonging to GO:0090150 using the Mashup-derived network embedding representations and (d) the STRING2GO_Mashup_ functional representations transformed by t-SNE.

We also display an example of the increased distance between two classes of proteins when predicting the term GO:0090150, which shows the highest improvement on the classifier training quality obtained by using STRING2GO_Mashup_+s_VM_, compared by using Mashup+SVM. Fig 5.b-5.c respectively show the 2-D visualization of raw Mashup-derived network embedding representations and the corresponding STRING2GO-learnt functional representations after transforming by t-SNE [41]. The red dots denote the protein samples belonging to class “Annotated”, while the green dots denote the protein samples belonging to class “Not-annotated”. The red dots are distributed in the similar scale of green dots in Fig 5.b, whereas the most of red dots are clustered in the right side in Fig 5.c. This fact indicates that the functional representations successfully encode higher discriminating power against two classes of protein samples.

## Conclusion

In this work, we present a novel deep learning-based protein function prediction method STRING2GO, which successfully learns a novel type of functional representations to train the down-stream classifiers for making predictions. STRING2GO shows the highest accuracy when predicting biological process protein functions, compared with other state-of-the-art network embedding representation-based protein function prediction methods. Based on this STRING2GO learning framework, there is potential for further improving the predictive accuracy by integrating representations from other data sources with the current PPI network embedding representations in a future study.

## Supporting information

**Table S1** List of 204 biological process Gene Ontology terms studied in this work.

**Table S2** Two-tailed Wilcoxon signed-rank tests at 0.05 of significance level on F1_GO_ scores obtained by different pairs of prediction methods over the hold-out evaluation.

**Table S3** Friedman test with Holm post-hoc correction results on F1_GO_ scores obtained by different prediction methods over the hold-out evaluation.

## Acknowledgements

The authors acknowledge the use of the high performance computing facility of the Department of Computer Science at University College London in the completion of this work.

